# Distinct Cellular Effects of Myotonic Dystrophy type 2 RAN Tetrapeptides in *Drosophila melanogaster*

**DOI:** 10.1101/2025.10.28.685039

**Authors:** Marta Marzullo, Assia De Simone, Marta Terribili, Michela Di Salvio, Degisew Yinur Mengistu, Maria Patrizia Somma, Rodrigo D’Amico, Gianluca Canettieri, Gianluca Cestra, Laura Ciapponi

**Author notes:** These authors contributed equally to the work (Marta Marzullo, Assia De Simone, Marta Terribili).

## Abstract

Myotonic dystrophy type 2 (DM2) is an autosomal dominant, multisystemic disorder caused by the expansion of CCTG repeats in the first intron of the CNBP gene. Repeat-associated non-AUG (RAN) translation of the expanded CCTG RNA may produce two tetrapeptide repeat proteins (TPRs), poly-QAGR and poly-PACL, whose roles in DM2 pathogenesis remain poorly understood. To investigate their individual contributions, we expressed codon-optimized QAGR and PACL peptides with ATG start codon in *Drosophila melanogaster*. Expression of QAGR and PACL TPRs significantly compromised fly viability and lifespan, induces eye degeneration and locomotor defects. We found that QAGR accumulated in the nucleolus, disrupted nucleolar integrity, and compromised rRNA processing. Moreover, QAGR expression interfered with autophagy, leading to the accumulation of Atg8a– and Ref(2)P-positive aggregates. Genetic interaction studies showed that overexpression of Atg8a or Ref(2)P mitigated QAGR-induced eye-toxicity, while knockdown of autophagy genes exacerbated it.

Conversely, PACL repeats promoted stress granule formation, as indicated by their colocalization with TIAR in human cells and their epistatic interaction with the *Drosophila* orthologue Rox8. Notably, cytoplasmic PACL aggregates were observed in myoblasts of DM2 patient. Together these findings demonstrate that QAGR and PACL peptides exert distinct toxic effects, impairing nucleolar function and autophagy, or altering stress granule dynamics, respectively. Both mechanisms likely converge to drive DM2 pathogenesis and represent promising therapeutic targets.

## INTRODUCTION

Myotonic dystrophy type 2 (DM2, OMIM 602668) is a multisystemic autosomal dominant disease that displays a wide spectrum of clinical manifestations, including proximal myotonia, degeneration of muscle fibres, defective cardiac conduction, cataracts, insulin resistance and other endocrine disorders (Meola, 2020; Meola and Cardani, 2015).

The genetic basis for DM2 is the expansion of an unstable CCTG repeat on chromosome 3q21, in the first intron of the cellular nucleic acid-binding protein (*CNBP*) gene, also named *ZNF9* (zinc finger protein 9; Liquori et al., 2001). The cause for the unstable expansion is unknown, however, the expanded DM2 alleles are strongly unstable with significant increase in length over time. The size of the (CCTG)n expansion is below 30 repeats in healthy individuals, whereas in DM2 patients is larger than 75 CCTG and can reach 11,000 repeats (Bachinski et al., 2009; Liquori et al., 2001).

Although the pathogenic mechanisms in DM2 are still not fully understood, several models have been proposed (Marzullo et al., 2023a). Accordingly, the CCTG expansion may affects CNBP expression *in cis* by altering transcription, impairing mRNA processing, or causing nuclear sequestration of the transcripts. Haploinsufficiency of *CNBP* gene has been suggested to play an important role in DM2 pathogenesis: indeed, mice carrying homozygous or heterozygous deletion of the CNBP allele develop clinical manifestations strongly reminiscent of DM2 myopathy (Wei et al., 2018), and downregulation of CNBP expression in *Drosophila* muscle tissues, causes severe locomotor defects rescued by reconstitution with either *Drosophila* or human CNBP (Coni et al., 2021). A gain-of-function RNA-mediated mechanism involves the transcription of CCTG repeats, resulting in the production of toxic CCUG-expanded RNA. This expanded RNA is associated with three primary pathological gain-of-function mechanisms: (i) formation of toxic RNA foci; (ii) general splicing defects, related to sequestration and dysregulation of RNA-binding proteins, such as the muscleblind-like proteins (MBNL1-3) and CUG-binding protein 1 (CUG-BP1) (Kanadia et al., 2006; Mohan et al., 2014); (iii) specific impaired CNBP pre-mRNA splicing, where the CCUG expansions potentially alters RNA structure and/or obstruct splicing regulators (Sznajder et al., 2018). Finally, non-ATG-mediated translation (RAN) have been also demonstrated (reviewed by Zu et al., 2018). Ectopic RAN translation has been reported in several degenerative diseases caused by microsatellite expansions such as SCA8, fragile X-associated tremor ataxia syndrome (FXTAS), C9ORF72-mediated amyotrophic lateral sclerosis/frontotemporal dementia (ALS/FTD), Fuchs endothelial corneal dystrophy, SCA31, Huntington disease, DM1 and DM2 (reviewed in Guo et al., 2022).

In particular, the expanded DM2-CCTG repeats are transcribed bidirectionally and the resulting RNAs undergo RAN translation, producing tetrapeptide repeat proteins of Pro-Ala-Cys-Leu (PACL) sequence from the sense strand and Gln-Ala-Gly-Arg (QAGR) tetrapeptide from the antisense strand. Both repeated tetrapeptides accumulate in DM2 patient brain and skin tissues and seem to be responsible for at least some of the neurological symptoms of DM2 patients (Rimoldi et al., 2025; Tusi et al., 2021; Zu et al., 2017).

Each of these potential mechanisms of toxicity is likely to contribute to disease initiation and progression, however, it is unclear to what extent each of them contributes to the development and to the clinical manifestations of the disease, and how they interact with each other (Malik et al., 2021). It has recently been proposed that at early stages CNBP haploinsufficiency and toxic nuclear RNAs are the main pathogenic mechanisms, while later the mRNAs accumulate in the cytoplasm where RAN translation occurs leading to production of toxic tetrapeptides, which may contribute to the worsening of the phenotype (Malik et al., 2021; Marzullo et al., 2023a).

Interestingly, the accumulation of different RAN-translated peptides in distinct cellular compartments may lead to unique pathogenic mechanisms and, consequently, variable disease presentations and severities in microsatellite expansion disorders. Although the precise role of RAN-translated peptides remains under investigation, their discovery has added a new layer of complexity to these diseases, with evidence suggesting that they are toxic, aggregating proteins contributing to disease progression.

Despite these insights, the molecular mechanisms underlying the exact contribution of DM2 RAN peptides to cellular dysfunction and pathogenesis remains largely unknown. To address this, we characterized the effects of PACL and QAGR tetrapeptides using *Drosophila melanogaster* as an *in vivo* model system. The two TPRs displayed distinct subcellular localizations and influenced separate cellular processes: QAGR accumulated in the nucleolus and modulated autophagy, whereas PACL localized predominantly in the cytoplasm and promoted stress granule formation. These findings suggest that RAN-translated tetrapeptides contribute to DM2 pathology through discrete, peptide-specific cellular mechanisms.

## RESULTS

### 1. Generation and characterization of DM2 TPRs transgenic flies

To analyse the specific contribution of the DM2 TPRs *in vivo* we have generated transgenic flies bearing a codon-optimized sequence that differs from the natural CCTG sequence and that encode either the PACL (TTG CCA GCT TGT) or the QAGR (CAG GCT GGA CGT) protein repeats (75 repetition each; Figure 1A). The coding sequences have been placed under the control of a UAS inducible element, after a canonical ATG starting codon and C-terminal tagged with a 3HA peptide (Figure 1A). Therefore, different GAL4 drivers were used to induce the expression of 75 repeats of PACL or QAGR HA-tagged tetrapeptides in different tissues.

**Figure 1.**
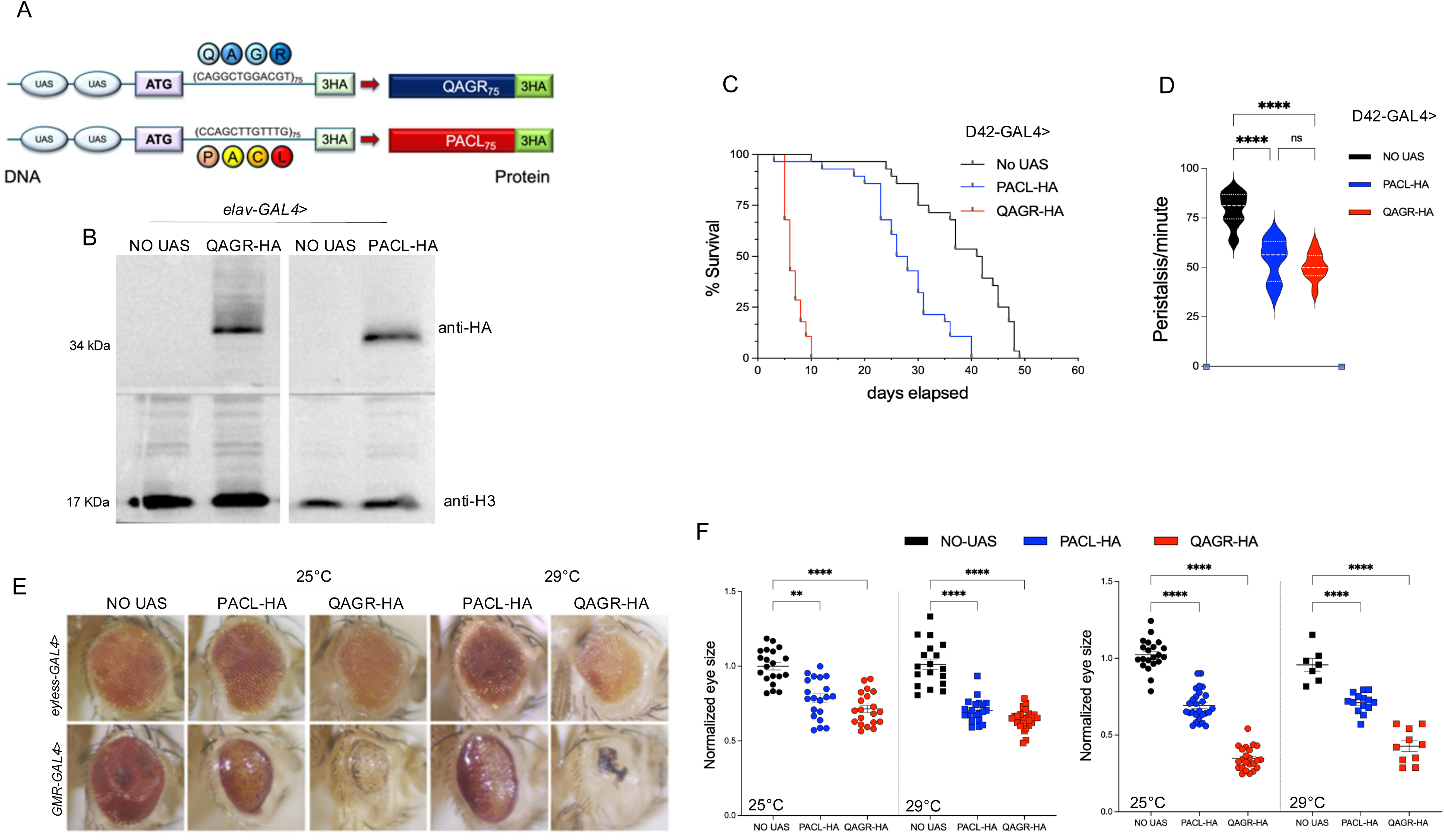
Expression of DM2 tetrapeptide repeats in motor neurons reduces lifespan, impairs locomotion and causes eye degeneration. (A) Schematic of transgenic “protein-only” constructs that have been generated using codons alternative to those found within the CCTG repeat arrays. Expansions were led by a canonical ATG codons, tagged with a triple HA epitope (3HA) and expressed under the control of the GAL4/UAS system. (B) Immunoblot of protein extracts obtained from *elav*-GAL4> UAS-PACL-3HA or UAS-QAGR-3HA larval brains, labelled with anti-HA antibody to detect the transgenic proteins and anti-H3 as loading control. The progeny of wild type individuals crossed to the driver (NO UAS) were used as control. (C) Survival rate as a function of time for flies expressing DM2 repeats under the control of the *D42*-GAL4 driver, shows a drastic reduction in lifespan of flies expressing UAS-QAGR-3HA (red line), and a milder yet noteworthy effect upon UAS-PACL-3HA expression (blue line), compared to controls (NO UAS, black line); n≥ 30 animals for each genotype. (D) Graphical representation of the distribution of peristaltic contractions performed in 1 min by third instar larvae expressing UAS-PACL-3HA (blue violin), UAS-QAGR-3HA (red violin), or NO UAS (black violin) under the control of *D42*-GAL4 driver at 25°C. **** P<0.0001, calculated by one-way ANOVA; error bars represent SEM with ≥ 15 larvae tested for each genotype in at least three independent experiments. (E) Representative adult eyes of flies expressing either UAS-PACL-3HA or UAS-QAGR-3HA under the control of the *GMR*-or *eyeless*-GAL4 driver. Flies were mated and reared at 25 °C or 29 °C, as indicated. (F) Statistical analysis of eye area of the same genotypes showed in E. The *Drosophila* eye arbitrary size was measured using FIJI. Each dot represents a single eye. **** P<0.0001; *** P<0.001, calculated by one-way ANOVA; error bars represent SEM with n≥ 20. As controls, progeny of wild-type individuals crossed to the used driver (NO UAS) were used.

To assess the expression efficacy of the transgenes, we performed immunoblotting analysis on protein extracts from larval brains expressing either UAS-(QAGR)_75_-3HA or UAS-(PACL)_75_-3HA (hereinafter referred to as UAS-PACL and UAS-QAGR) under the pan neuronal driver *elav*-GAL4. The results obtained showed that both TPRs are significantly expressed (Figure 1B). Therefore, this transgenic fly model is well-suited to investigate the specific cellular and molecular mechanisms of the TPRs-based toxicity and its role in DM2 pathogenesis.

### 2. DM2 TPRs reduce lifespan, impair locomotion and induce eye degeneration

To address the toxic effect of DM2 tetrapeptides in cells involved in the control of muscle contraction, we analyzed the phenotypic consequences of PACL and QAGR repeats expression specifically within motor neurons, using the *D42*-GAL4 driver. Flies expressing QAGR TPRs in motor neurons exhibited a significant reduction in their lifespan with a median survival time of 6 days for QAGR compared to 41.5 days for wild type controls. The impact of PACL repeats was less pronounced, but still significant compared to controls, with an average lifespan of 27 days (Figure 1C). Interestingly, TPRs toxicity also affected larval locomotor capabilities. Larval locomotion is characterized by peristaltic waves of muscle contractions that propagate along the body through coordinated segmental movements. Expression of either PACL or QAGR repeats in motor neurons significantly reduced peristaltic movements compared to controls (Figure 1D).

*Drosophila* eye provides an ideal tissue for studying the genetic regulation of neurodegeneration and allow to study the effect of the expression of toxic products. Moreover, eye expression of pure, uninterrupted CCUG-repeat expansions has been shown to cause strong retina degeneration (Mishra and Knust, 2013; Yu et al., 2015). Thus, we investigated whether the expression of PACL and QAGR DM2 TPRs early (*ey*-GAL4) or late (*GMR*-GAL4) during eye development causes retina degeneration. Expression of the UAS-(PACL)_75_-3HA or UAS-(QAGR)_75_-3HA construct under the control of eye-specific drivers led to a significant reduction in eye area, abnormal pigmentation, and a smooth eye surface, indicative of neurodegeneration and strongly resemble the phenotypes observed in previous fly models of DM2 (Figure 1E and F) (Yenigun et al., 2017; Yu et al., 2015). Immunofluorescence analysis on larval brains cells expressing the transgenes under the pan neuronal driver *elav*-GAL4, showed that PACL TPRs were predominantly cytoplasmic, exhibiting both diffuse and punctate staining. Conversely, QAGR repeats displayed a distinct nuclear pattern (Figure 2A), with a specific staining around the nucleolus, as indicated by its proximity with the well-established nucleolar marker fibrillarin (Figure 2A). Similar observations were obtained in brain cells from larvae expressing the transgenes under the motor neuronal driver *D42*-GAL4.

**Figure 2.**
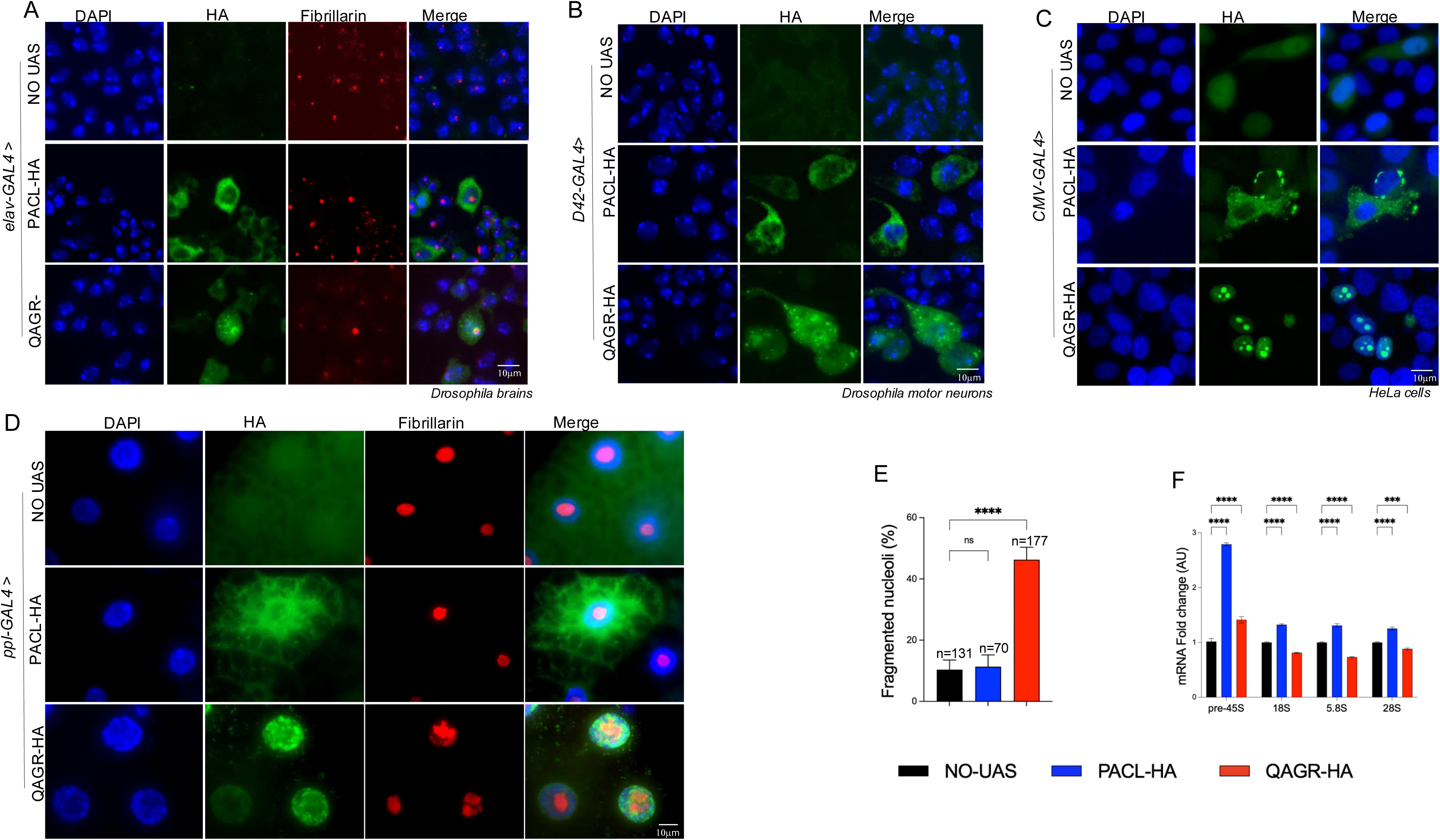
QAGR TPRs expression induces nucleolar stress. (A) Representative images of *Drosophila* larval brain cells from controls (NO UAS) or from larvae expressing either UAS-PACL-3HA or UAS-QAGR-3HA construct, under the control of the *elav*-GAL4 driver. Cells were stained with DAPI (blue), anti-HA (green) and anti-fibrillarin (red) antibodies. Note the distinct cellular expression patterns between the two tetrapeptides, with PACL leading to a primarily cytoplasmic diffuse localization and QAGR forming aggregates around the nucleolus (Scale bar: 10 µm). (B) Representative images of *Drosophila* larval motor neurons from controls (NO UAS) or from larvae expressing either UAS-PACL-3HA or UAS-QAGR-3HA tetrapeptide repeats, under the control of *D42*-GAL4. Cells were stained with DAPI (blue) and anti-HA antibody (red). (Scale bar: 10 µm). As controls, progeny of wild-type individuals crossed to the driver (NO UAS) were always used (C) Representative images of HeLa cells transfected with either UAS-PACL-3HA or UAS-QAGR-3HA plasmid together with the *pCMV*-GAL4 plasmid (scale bar:10 µm), stained with DAPI (blue) and anti-HA antibody (green). Cell transfected only with *pCMV*-GAL4 plasmid were used as control (NO UAS). Note that PACL exclusively localizes to the cytoplasm, while QAGR is specifically distributed in the nucleolus. (D) Representative larval fat bodies from individual expressing either UAS-PACL-3HA or UAS-QAGR-3HA construct, under the control of the *ppl*-GAL4 driver stained with anti-HA (green) and fibrillarin (red) antibodies. (E) Quantification of percentage of fragmented nucleoli in fat body’s cells as in A, n represents the total number of nuclei scored. **** P<0.0001, calculated by one-way ANOVA; error bars represent SEM. (F) Quantitative PCR on total RNA extracts from UAS-PACL-3HA or UAS-QAGR-3HA expressing larvae under the control of the *ppl*-GAL4 driver. **** P<0.0001, *** P<0.001 calculated by one-way ANOVA. Error bars represent SEM with n=2 for biological replicates and n=6 for technical replicates. As controls, progeny of wild-type individuals crossed to the driver (without UAS) were used.

PACL repeats had diffuse and specific cytoplasmic localization with the formation of rare cytoplasmic aggregates, while QAGR repeats form aggregates localized in both cytoplasm and nucleus (Figure 2B). Analogous distinct subcellular localization patterns were observed in HeLa cells (Figure 2C), consistent with previous observations in transfected HEK293T cells (Zu et al., 2017).

### 3. QAGR repeats induce nucleolar stress and affect autophagy

In the *Drosophila* fat bodies, nucleolar defects can have systemic effects, including an indirect impact on muscle function, likely due to altered protein synthesis, energy metabolism, and stress signalling. Further, as fat body cells are highly engaged in metabolic and immune functions, their nucleoli are typically well-developed to support high ribosome production. Immunofluorescence analysis of fat body cells expressing UAS-QAGR under the control of the specific driver *ppl*-GAL4 confirmed that QAGR TPRs localize both in nucleus and cytoplasm, in line with the phenotype observed in larval brains. Specifically, we confirmed the peri nucleolar localization of QAGR as ascertained by co-labelling with the nucleolar marker fibrillarin (Figure 2D). Furthermore, expression of QAGR led to altered nucleolar morphology and evident nucleolar fragmentation, (Figure 2D and E). Since, perturbation of nucleolar morphology could ultimately impair ribosomal RNA biogenesis, we measured the levels of 5,8S, 18S and 28S rRNA, which are processed from the precursor 45S rRNA. qRT-PCR analysis revealed a significant decrease in the levels of mature 5.8S, 18S, and 28S rRNAs in larvae expressing QAGR-TPRs, accompanied by a marked accumulation of pre-rRNA 45S compared to controls (Figure 2F). These results suggest that QAGR-TPRs expression affect pre-rRNA processing and maturation, impairing nucleolar function, as suggested by the observed nucleolar fragmentation.

Immunofluorescence analysis of fat body cells expressing UAS-PACL under the control of *ppl*-GAL4 confirmed the cytoplasmatic localization observed in larval brains. Notably, PACL TPRs had no detectable effect on nucleolar morphology but caused a general upregulation of both pre-rRNA 45S and mature rRNAs (Figure 2D, E and F), suggesting that PACL expression induces a stress response that enhances rDNA transcription and rRNA processing without perturbing nucleolar integrity.

To assess the effects of the DM2 TPRs in muscle, the most affected tissue in DM2 patients, we expressed PACL or QAGR tetrapeptide repeats under the control of the muscle-specific *Mef2-* GAL4 driver. Notably, expression of both TPRs in muscle led to adult lethality at 25°C. Lowering the temperature to 18°C enabled to obtain QAGR-expressing third instar larvae, whereas PACL expression remained lethal even under these conditions.

As shown in Figure 3A, larvae expressing QAGR TPRs in muscles exhibit severe locomotor defects, evidenced by a significant reduction in peristaltic wave frequency. These defects associate with the accumulation of QAGR TPR aggregates in muscle tissue, as revealed by multiple HA-positive inclusions detected through immunofluorescence in larval muscle fillets (Figure 3B and C). Because the accumulation of toxic peptides is often associated with autophagy, to elucidate how QAGR TPR aggregation contributes to larval locomotor defects, we investigated whether these peptides affect autophagy, a process that plays a crucial role in the clearance of toxic protein aggregates and damaged organelles (Boivin et al., 2020; Ciechanover and Kwon, 2015).

**Figure 3.**
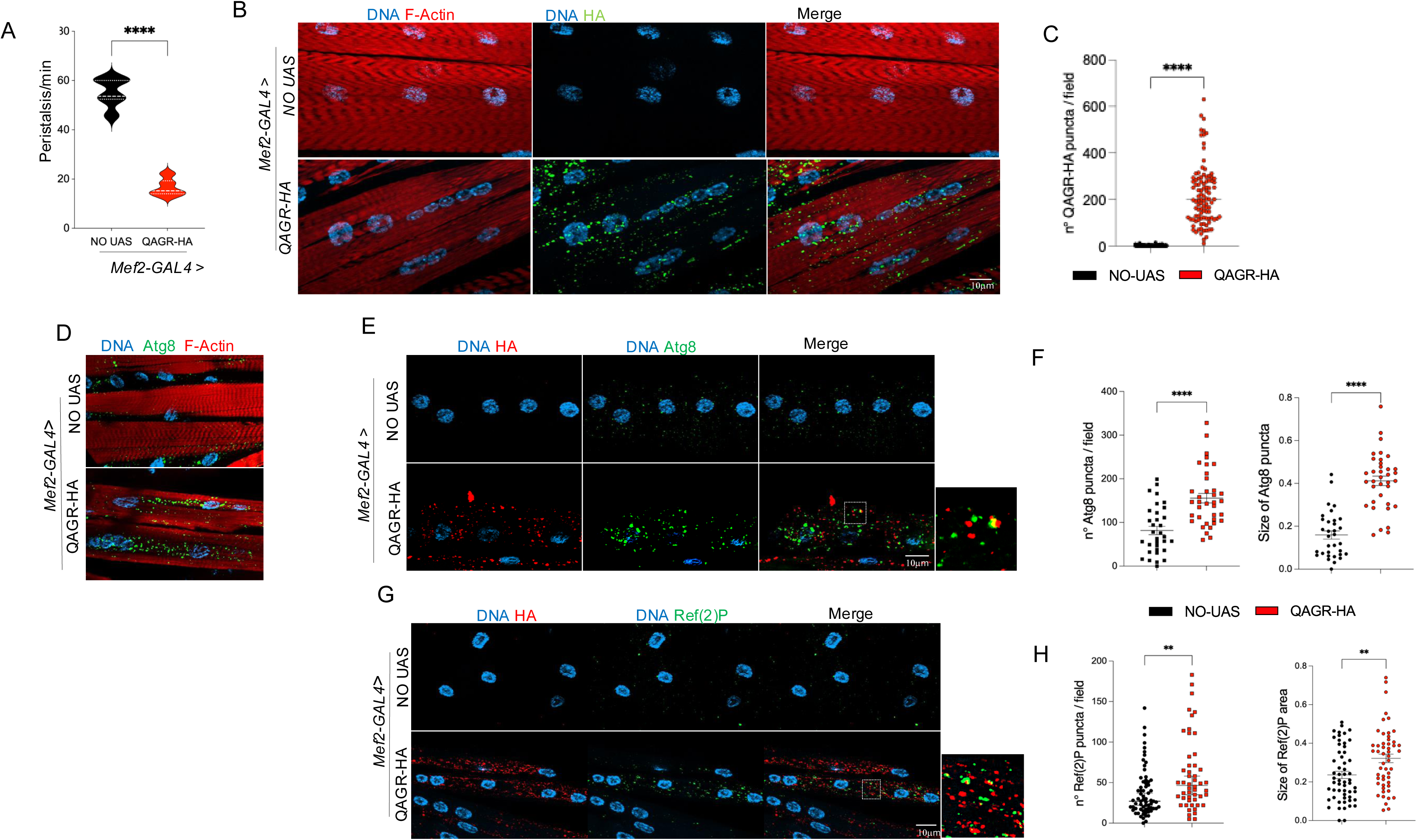
QAGR-TPRs form aggregates in larval muscles, affecting locomotor activity and autophagy. (A) Graphical representation of the number of peristaltic contractions performed in 1 min by control larvae (NO UAS; black violin) or UAS-QAGR-3HA expressing larvae (red violin). **** P<0.0001, calculated by t-test; error bars represent SEM with ≥ 10 larvae tested for each genotype. The low number of larvae used for the experiment is justified by the semi-lethality of the informative progeny. (B) Representative confocal images of third instar larval muscle fibres expressing or not (NO UAS) UAS-QAGR-3HA under the control of the muscle-specific driver *Mef2-*GAL4 at 18°C stained with anti-HA (green) and phalloidin, a marker for F-actin (red), antibodies, and DAPI (DNA in blue). Scale bar 10 μm. (C) Graphical representation of the number of QAGR puncta/field is shown (rounds) present in control larval muscles (NO UAS; black) and in UAS-QAGR-3HA expressing larval muscle (red). **** P<0.0001, calculated by t-test; error bars represent SEM with ≥ 100 fields tested for each genotype. As controls, progeny of wild-type individuals crossed to the driver (NO UAS) were used. (D) Representative confocal images of third instar larval muscle fibres expressing, or not (NO UAS), UAS-QAGR-HA under the control of the muscle-specific driver *Mef2-*GAL4 at 18°C. Larval muscles were stained with fluorescent labelled phalloidin, a marker for F-actin (in red), anti-Atg8a antibody (in green) and DAPI (in blue). Note that the QAGR-expressing muscle fibres display a high number of Atg8a positive puncta. (Scale bar: 10 mm). (E) Representative confocal images of third instar larval muscle fibres expressing, or not (NO UAS), UAS-QAGR-3HA under the control of the muscle-specific driver *Mef2-*GAL4 at 18°C. Larval muscles were stained with anti-HA (red) and anti-Atg8a (green) antibodies and DAPI (DNA, blue). (Scale bar: 10 μm). Dashed square represents the area shown in high magnification on the right. (F) Graphical representation of the number (squares) and size (circles) of Atg8a puncta present in control larval muscles (NO UAS; black) and in UAS-QAGR-3HA expressing larval muscle (red). ****P<0.0001, calculated by t-test; error bars represent SEM with ≥ 30 fields tested for each genotype. (G) Representative confocal images of third instar larval muscle fibres expressing or not (NO UAS), UAS-QAGR-3HA under the control of the muscle-specific driver *Mef2.*GAL4 at 18°C. Larval muscles were stained with anti-HA (red) and anti-Ref(2)P (green) antibodies, and DAPI (DNA, blue). (Scale bar: 10 μm). (H) Graphical representation of the number (squares) and size (circles) of Ref(2)P puncta present in UAS-QAGR-3HA expressing larval muscle (red) compared to controls (NO UAS, black). **** P<0.0001, **P<0.001calculated by t-test; error bars represent SEM with ≥ 50 fields tested for each genotype.

The microtubule-associated protein light chain 3 variants (LC3s) and their paralogs GABARAPs are cleaved, processed, and inserted into nascent autophagosomes, where they participate in autophagosome formation and cargo selection for degradation (Lamark and Johansen, 2021). In *Drosophila* the Autophagy-related gene 8a (Atg8a) protein serves as the functional homolog of both human LC3 and GABARAP and is widely used as a marker of autophagic vesicles.

To study autophagosome activity upon TPRs expression, we performed immunostaining of larval muscles expressing QAGR using an antibody that labels fly Atg8a/GABARAP. This analysis revealed a marked increase in the number of Atg8a-immunoreactive puncta in larval muscles expressing UAS-QAGR (*Mef2*-GAL4) compared to controls (Figures 3D, E and F). This phenotype closely resembles that observed under starvation conditions, suggesting that QAGR expression may activate the autophagy pathway, mimicking a nutrient-deprivation phenotype (Allen et al., 2020; Érdi et al., 2012; Scott et al., 2004; Zirin et al., 2015). Immunofluorescence staining for Ref(2)P, the fly homolog of the selective autophagy receptor p62 (Zatloukal et al., 2002), revealed also a significant increase in Ref(2)P-positive puncta in QAGR-expressing muscles compared with controls (Figures 3G and H). Notably, both Atg8a– and Ref(2)P-positive puncta were not only more numerous but also increased in size following QAGR expression (Figures 3E-H). Together, these findings indicate that QAGR expression challenges autophagy, causing both accumulation and enlargement of autophagic vesicles. According with this hypothesis, Ref(2)P overexpression partially rescue the QAGR-dependent eye degeneration (Figure 4A and B). Further, RNAi-mediated knockdown of various autophagic genes (Atg5, Atg7, and Atg8; Mizushima et al., 2011), specifically exacerbated QAGR TPRs-induced eye degeneration (Figures 1, and Figure 4C and D). Interestingly, overexpression of Atg8a significantly ameliorated QAGR TPRs-induced eye degeneration (Figure 4E and F). All together these data suggest that although QAGR expression activates autophagy, its potentiation mitigates QAGR-mediated toxicity.

**Figure 4.**
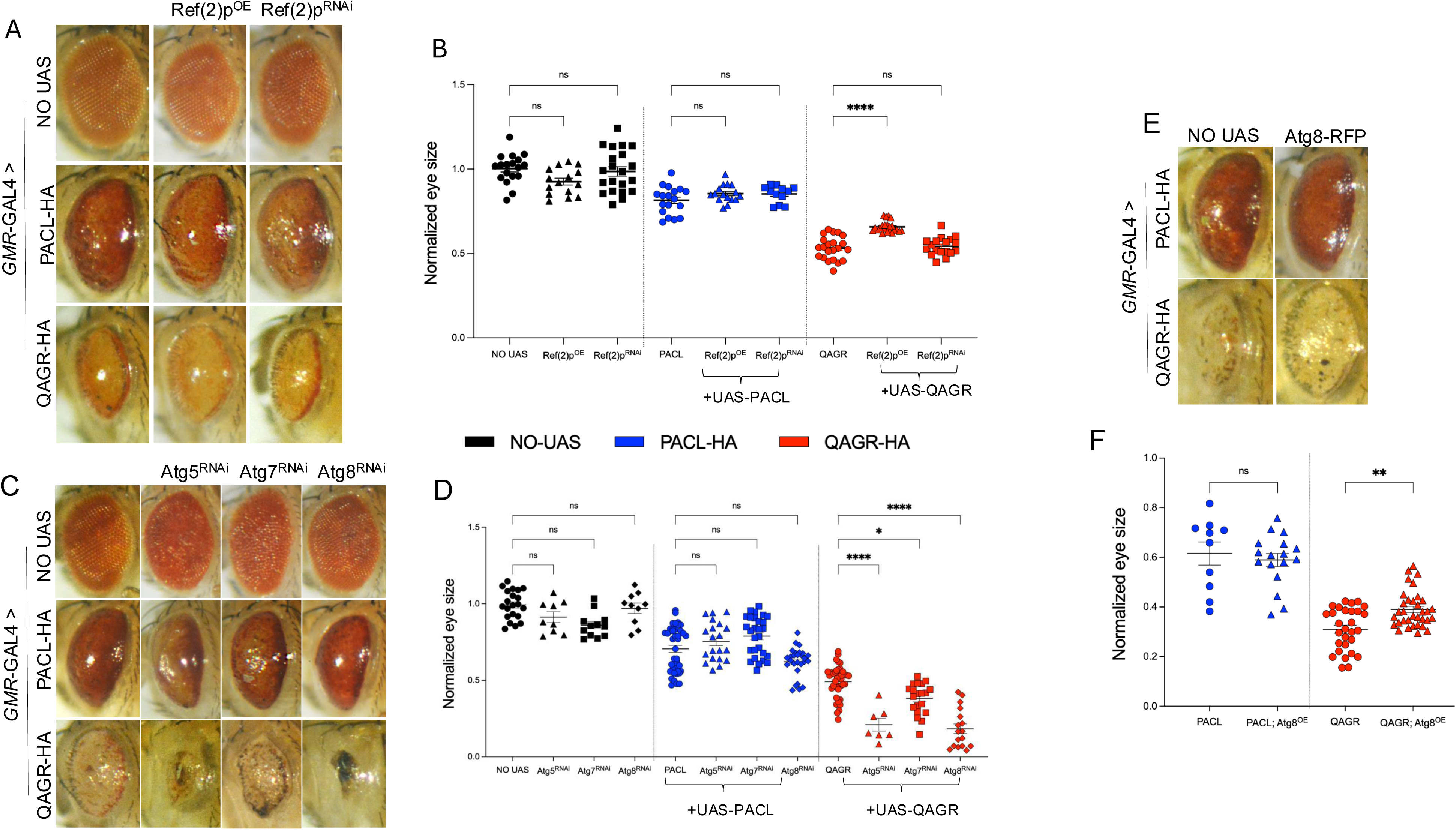
QAGR-dependent eye degeneration is influenced by autophagic genes modulation. (A) Representative adult eyes of flies expressing, or not (NO UAS), either UAS-PACL-3HA or UAS-QAGR-3HA under the control of the *GMR*-GAL4 driver, alone or in combination with either Ref(2)P silencing (UAS-Ref(2)P^RNAi^) or overexpression (UAS-Ref(2)P^OE^). Flies were mated and reared at 25 °C. (B) Graphical representation of the mean eye size of the same genotypes showed in A. The *Drosophila* eye size was measured by using FIJI. **** P<0.0001, not significative, calculated by one-way ANOVA; error bars represent SEM with n≥ 15. As controls, progeny of wild-type individuals crossed to the driver (NO UAS) were used. (C) Representative adult eyes of flies expressing, or not (NO UAS), either UAS-PACL-3HA or UAS-QAGR-3HA under the control of the *GMR*-GAL4 driver, alone or in combination with either Atg5 (UAS-Atg5^RNAi^), Atg7 (UAS-Atg7^RNAi^) or Atg8a (UAS-Atg8^RNAi^) silencing. Flies were mated and reared at 25 °C. (D) Graphical representation of the mean eye size of the same genotypes showed in A. **** P<0.0001, ** P<0.01, * P<0.1, ns, not significative, calculated by one-way ANOVA; error bars represent SEM with n≥ 20. (E) Representative adult eyes of flies expressing, or not (NO UAS), either UAS-PACL-3HA or UAS-QAGR-3HA under the control of the *GMR*-GAL4 driver, alone or in combination with Atg8 overexpression (UAS-Atg8^OE^). Flies were mated and reared at 25 °C. (F) Graphical representation of the mean eye size of the same genotypes showed in E. ** P<0.01, not significative, calculated by one-way ANOVA; error bars represent SEM with n≥ 20. The *Drosophila* eye size was measured by using FIJI. As controls, progeny of wild-type individuals crossed to the driver (NO UAS) were used.

### 4. PACL TPRs associate with stress granules

Our observations on fly eye neurodegeneration suggest that PACL-mediated toxicity is not affected by modulation of the autophagy factors (Figure 4). Interestingly, PACL TPRs primarily localized in the cytoplasm, where they occasionally formed granules (Figure 2). Dipeptide repeat proteins (DPRs) such as poly-GA, poly-GP, and poly-PA, generated by RAN translation from the expanded *CCCGGG* repeat in the *C9orf72* gene, have been shown to induce stress granule (SG) formation (Tao et al., 2015). Thus, we assessed whether PACL TPRs contribute to SG formation. To determine whether PACL-containing granules colocalized with SGs, we performed co-immunofluorescence with antibodies against TIAR, an RNA-binding protein essential for SG assembly. To promote stress granule formation, we treated cells with sodium arsenide, which induces prominent oxidative stress that activates eIF2α phosphorylation, leading to global translational arrest (Jain et al., 2016). In untreated cells TIAR typically localizes in the nucleus, whereas after sodium arsenide treatment, it translocates to the cytoplasm where it promotes SG formation. Interestingly, in cells expressing PACL, TIAR-positive cytoplasmic SGs colocalized with PACL even in the absence of arsenide treatment (Figure 5A). This suggests a link between PACL repeats expression and SG formation, which may contribute to DM2 pathogenesis.

**Figure 5.**
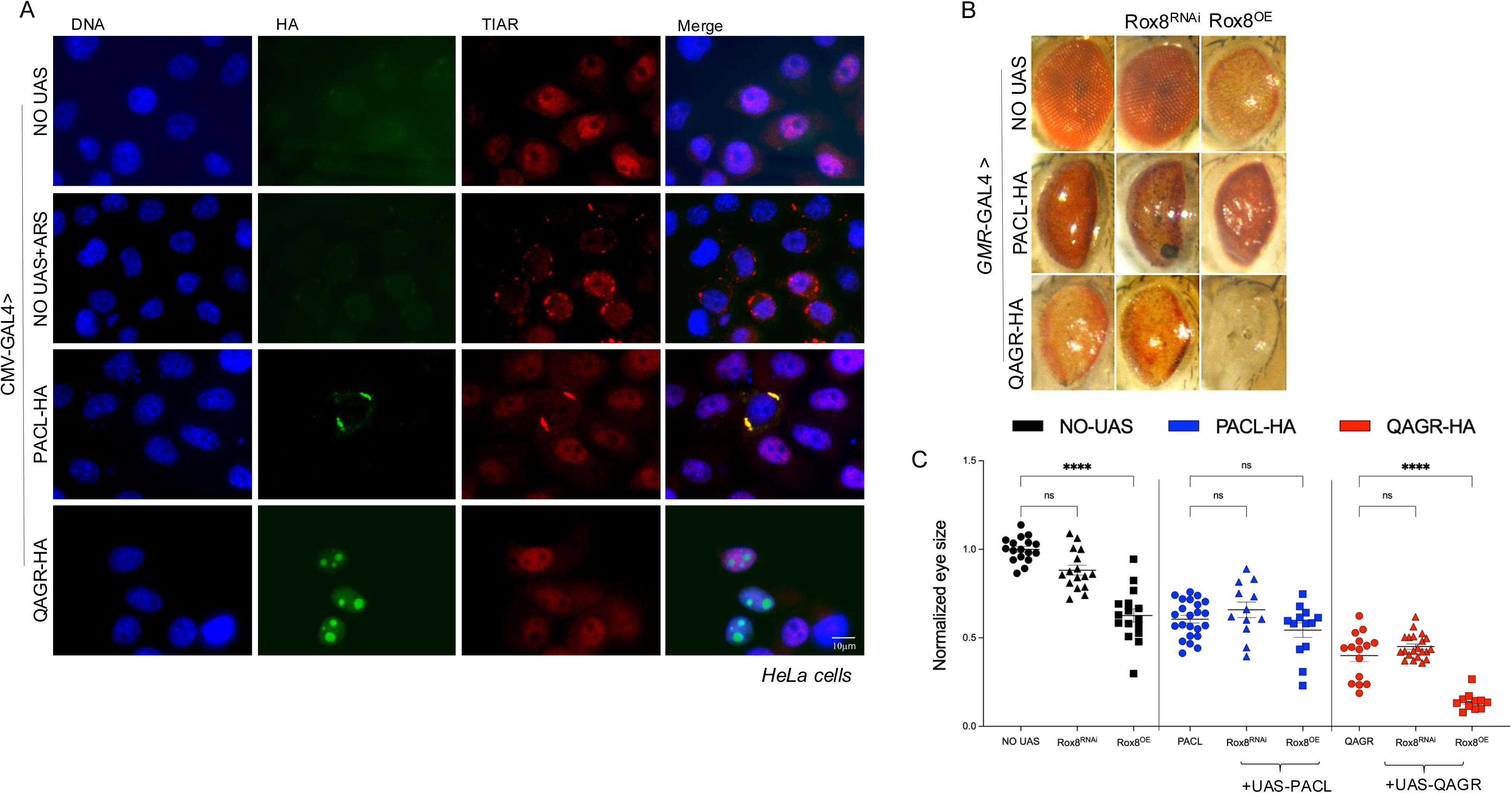
PACL TPRs colocalize with TIAR in HeLa cells. (A) Representative images of HeLa cells transfected with either UAS-PACL-3HA or UAS-QAGR-3HA plasmid together with the *pCMV*-GAL4 plasmid stained with anti-HA (green) and anti-TIAR antibodies (red), and DAPI (DNA, blue). HeLa transfected only with the *pCMV*-GAL4 plasmid treated or not with arsenide, were used as positive and negative control, respectively (scale bar:10 µm). (B) Representative adult eyes of flies expressing, or not (NO UAS), either UAS-PACL-3HA or UAS-QAGR-3HA under the control of the *GMR*-GAL4 driver, alone or in combination with either UAS-Rox8^RNAi^ or UAS-Rox8^OE^. Flies were mated and reared at 25 °C. (C) Graphical representation of the mean eye size of the same genotypes showed in B. \**\*\*\** P<0.0001, *\** P<0.1, ns, not significative, calculated by one-way ANOVA; error bars represent SEM with n≥ 20. The *Drosophila* eye size was measured by using FIJI. As controls, progeny of wild-type individuals crossed to the driver (NO UAS) were used.

Conversely, no SG activation was observed upon QAGR peptide expression (Figure 5A). Genetic interaction experiments with Rox8, the *Drosophila* orthologue of human TIAR, showed that changing Rox8 levels (either by silencing or overexpression) did not affect PACL-induced eye degeneration. This strongly suggests an epistatic relationship between PACL and TIAR in the SG formation pathway, in which PACL-mediated toxicity is not modified by any alteration of TIAR levels. In contrast, Rox8 overexpression, which *per sé* causes eye neurodegeneration, exacerbated the eye degeneration induced by QAGR TPRs, suggesting that alterations of SG dynamics, may have additive effects on the autophagy-related toxicity of QAGR (Figure 5B and C).

### 5. PACL TPRs accumulate in the cytoplasm of human DM2 myoblasts

To determine whether DM2 RAN proteins are expressed *in vivo* in muscle cells of DM2 patients, we used commercial polyclonal antibodies targeting the DM2 PACL and QAGR repeat motifs in *Drosophila* protein extracts. Immunoblot analysis showed that, while the anti-PACL antibody recognized transgenic 3HA-tagged proteins, the anti-QAGR antibody did not label them (Figure 6A). A similar pattern was observed in protein extracts from human myoblast of healthy individuals or from DM2 patient, where specific bands were detected for PACL peptides, but not for QAGR (Figure 6B).

**Figure 6.**
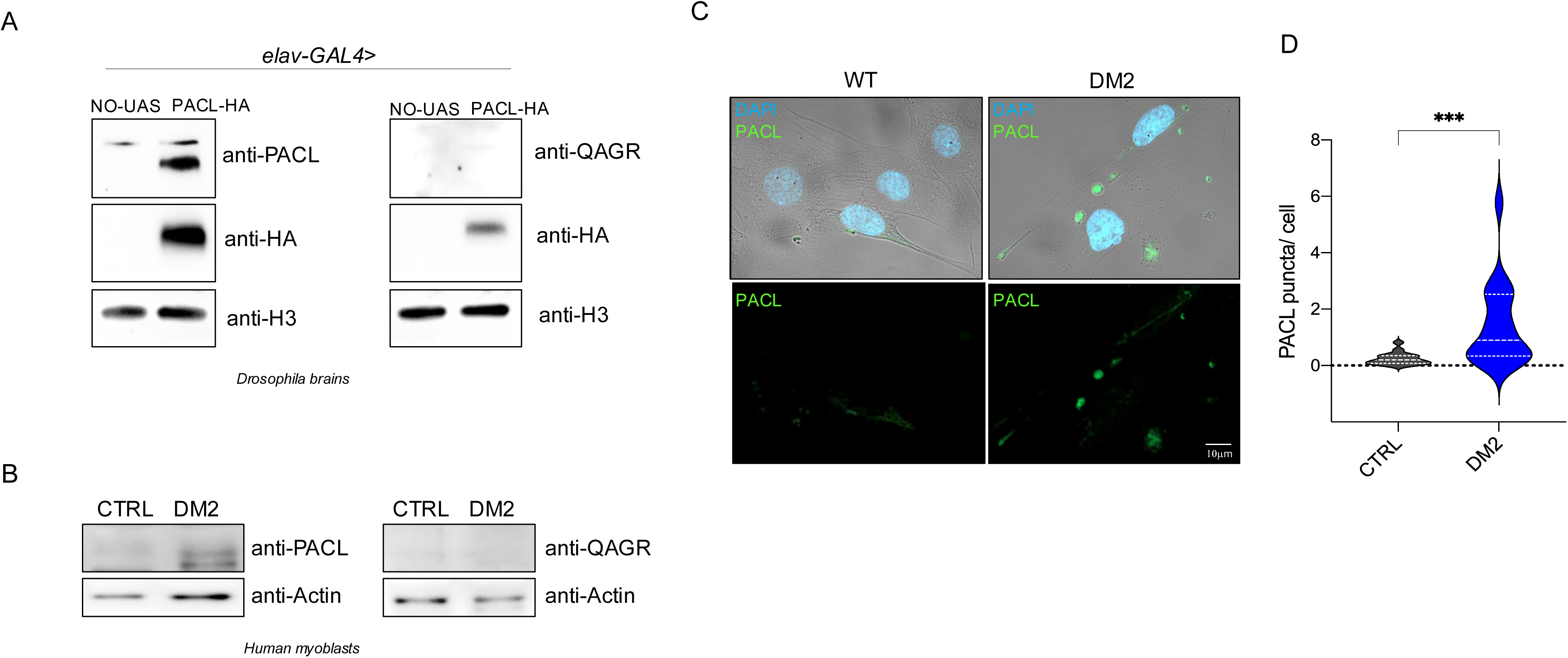
PACL TPRs accumulate in the cytoplasm of human DM2 myoblasts. (A) Immunoblot of *Drosophila* protein extracts from transgenic larval brains expressing either PACL-HA or QAGR-HA TPRs constructs under the control of the *elav*-GAL4 driver, labelled with anti-QAGR or anti-PACL antibodies. Anti vibrator was used as loading control. (B) Immunoblot of myoblasts protein extracts from control individuals and DM2 patients, labelled with anti-QAGR or anti-PACL antibodies. Actin was used as loading control. (C) Representative images human myoblasts from control individuals and DM2 patients (scale bar:10 µm), stained with DAPI (blue) and anti-PACL antibody (green). (D) Graphical representation of the number of PACL positive puncta observed in WT controls or DM2 human myoblasts, as shown in C. \**\*\** P<0.001, calculated by t-test with n≥ 50.

The DM2-RAN proteins were previously shown to accumulate in DM2 patient brains and in DM2 patients skin biopsies (Rösing et al., 2024; Zu et al., 2017), thus we tested DM2 expression in DM2 patient myoblast. Immunofluorescence analysis further confirmed the absence of a detectable QAGR signal in DM2 patient myoblasts (not shown), whereas staining with the anti-PACL antibody revealed the formation of specific cytoplasmic aggregates, significantly higher in DM2 cells in both number and size with respect to wild type control myoblasts (Figure 6C and D).

## DISCUSSION

The use of codon-optimized *Drosophila* model expressing PACL and QAGR tetrapeptide repeats offers an incisive tool to dissect the protein-specific toxicity of DM2 RAN translation products. Crucially, the ‘protein-only’ strategy circumvents sequence-specific RNA-mediated effects, enabling a direct analysis of pathological mechanisms driven by repeat-encoded peptides. Thus, our study provides significant insights into the contribution of DM2 RAN-translated tetrapeptide repeat proteins to disease pathology.

A major finding is the distinct subcellular localization and aggregation tendencies of PACL and QAGR TPRs. While PACL repeats primarily exhibited a cytoplasmic distribution with occasional aggregates, QAGR repeats formed prominent aggregates in both cytoplasm and nucleus, with a marked propensity to accumulate within the nucleus. This may be due to the high density of repetitive arginine residues that enhance cell permeability and nuclear import by mimicking the positive charge typical of nuclear localization signals (Kwon et al., 2014).The nuclear localization of QAGR TPRs, particularly their close proximity with the nucleolus, suggests an effect on the nucleolar function. This is further supported by the alterations observed in the nucleolar morphology, as well as a reduction in 18S and 28S rRNA levels upon QAGR expression.

Conversely, the expression of PACL TPRs causes a significant upregulation of unprocessed pre-45S rRNA. Since nucleolar accumulation of unprocessed pre-45S rRNA is a well-known cellular stress response associated with the inhibition of global mRNA translation (Szaflarski et al., 2022), this PACL-mediated effect on ribosomal biogenesis is in agreement with its possible role on stress granule formation. Accordingly, PACL TPRs colocalize with stress granule components under basal conditions. The ability of PACL to induce SG formation even in the absence of oxidative stress indicates an intrinsic capacity to alter RNA metabolism and translational regulation, potentially via aberrant recruitment of RNA-binding proteins like TIAR.

A further finding of our work is the specific effect of QAGR expression in muscle. QAGR expression in muscle robustly upregulates autophagic proteins, as evidenced by increased Atg8a and Ref(2)P puncta formation. These findings suggest that the accumulation of QAGR TPRs could upregulate the autophagy pathway as a protective mechanism to clear toxic peptides. However, despite this activation, a substantial amount of toxic TPRs remains within the cell, contributing to phenotypic abnormalities observed in both muscle and eye tissues. Conceivably, Atg8a or Ref(2)P overexpression by enhancing QAGR TPR clearance, partially rescues the eye phenotype.

Conversely, silencing *Atg5*, *Atg7*, *Atg8* or *Ref(2)P* by disrupting autophagy, leads to increased retention of QAGR TPRs and enhances eye degeneration. Accumulation of repeated peptide with consequent activation of autophagy has been reported in several diseases caused by microsatellite expansions such as fragile X-associated tremor ataxia syndrome (FXTAS), C9ORF72-mediated amyotrophic lateral sclerosis/frontotemporal dementia (ALS/FTD) and Huntington disease (Alvarez-Mora et al., 2023; Ciechanover and Kwon, 2015; Mengistu et al., 2025; Pircs et al., 2018). Upregulating autophagy aims promote the clearance of toxic protein aggregates that contribute to the loss of neuron or muscle homeostasis. Although autophagy regulation is complex, its activation helps maintain cellular balance disrupted by pathogenic proteins. The selective modulation of specific steps of the autophagy process, such as autophagosome formation, cargo recognition, or fusion with lysosomes, can represent a promising strategy to reduce aggregates-mediated toxicity and slow disease progression.

In contrast, PACL TPRs show minimal impact on autophagy markers but colocalize with stress granule components under basal conditions. While PACL-mediated toxicity is not affected by modulation of autophagy genes, its effect on stress granule dynamics may still be relevant in contexts of cellular stress or aging, where SG misregulation contributes to disease pathogenesis. The absence of effects of modulated Rox8 levels on PACL-mediated neurodegeneration in the fly eye indicate that PACL TPRs act epistatically to Rox8. However, the association of PACL peptides with stress granules is particularly intriguing, as SG formation is a common cellular response to stress and has been implicated in several neurodegenerative diseases.

The analysis of human DM2 patient myoblasts further supports the relevance of our *Drosophila* model. Immunoblot and immunofluorescence analyses confirmed the presence of PACL peptides in DM2 patient-derived myoblasts, where they formed cytoplasmic aggregates. This aligns with our *Drosophila* findings, where PACL localized in the cytoplasm. Unfortunately, QAGR peptides were undetectable, because of antibody inefficacy.

In conclusion, our study highlights the distinct toxic contributions of PACL and QAGR TPRs to DM2 pathology. The differential effects of these peptides on nucleolar function, autophagy, and stress granule dynamics highlight the complexity of DM2 pathogenesis and offer potential avenues for therapeutic intervention. Future studies should focus on elucidating the precise molecular mechanisms underlying these toxic effects and exploring targeted strategies to counteract them in DM2 patients.

## Material and Methods

### *Drosophila* strains and rearing conditions

*Drosophila* stocks were maintained on standard fly food (25 g/L corn flour, 5 g/L lyophilized agar, 50 g/L sugar, 50 g/L fresh yeast, 2,5 ml/L Tegosept [10% in ethanol], and 2,5 ml/L propionic acid) at 25°C in a 12-hr light/dark cycle. All experiments were performed in the same standard conditions, at the temperature reported in figure legends.

Protein-only alternative codon constructs were synthesized by GeneArt, (Life Technologies). The plasmids for inducible expression of *UAS-*(*PACL*)_75_*-3HA* or UAS-(*QAGR*)_75_*-3HA* were generated by cloning the 3HA epitope CDS fused in-frame with the (CCAGCTTGTTTG)_75_ for PACL or (CAGGCTGGACGT)_75_ for QAGR into the UAS-attB vector, (GeneArt, Life Technologies). The *UAS-(PACL*)_75_*-3HA* or UAS-(*QAGR*)_75_*-3HA* plasmids were injected in y^1^ w^67c23^; P{CaryP}attP2 embryos (BDSC Stock #24749); germline transformation was performed by Bestgene Inc. (Chino Hills, California) using standard procedures. All the driver lines used have been previously described and available from the Bloomington stock centre.

### *Drosophila* larval locomotion analyses

Larval locomotor activity was measured by counting the number of peristaltic contractions of third instar larvae performed within 1 min on the surface of a 1% agarose gel in a Petri dish; measurements were repeated five times for each larva, at least 10 larvae per genotype in each experiment (Coni et al., 2021).

### Survival curves

Control and experimental flies were collected within 24 hours post eclosion, sorted by sex under carbon dioxide (CO_2_) anaesthesia and maintained in a climate chamber on a standard fly food at constant temperature (25 °C). Flies were kept 10 individuals per vial in triplicates. Dead flies were recorded daily, and fresh medium vials were provided two times per week. GraphPad Prism 6 through the Kaplan-Meier method was used for the survival analysis, by constructing a survival curve that tracks the survival rate over time.

### Immunoblot and antibodies

*Drosophila*: Protein extracts were derived from third instar larvae lysed in sample buffer, fractionated by SDS-PAGE and transferred to nitrocellulose membrane. Primary and secondary antibodies were diluted in 5% milk/ PBS-Tween 0,1% (GE Health Care).

*Human cells:* Cells were lysed in denaturing buffer SDS-urea (50 mM Tris HCl, pH 7.8, 2% sodium dodecyl sulphate (SDS), 10% glycerol, 10 mM Na4P2O7, 100 mM NaF, 6 M urea, 10 mM EDTA). Protein extracts were then sonicated, quantified, and resolved via SDS-PAGE before transfer to a nitrocellulose membrane (#NBA085C001EA, Perkin Elmer, Waltham, MA, USA). Primary and secondary antibodies were diluted in 5% milk/ PBS-Tween 0,1% (GE Health Care). Detection was performed by using WesternBright ECL (K-12045-D50, Advansta). Signals were detected and acquired using the Gel Doc II scanning system (BioRad). Bands densitometric analysis was performed using the ImageJ software1.50i. Primary antisera specifics and dilution used are reported in the resources table.

### HeLa cell transfection and immunofluorescence

HeLa cells were grown in Dulbecco’s Minimal Essential Medium (DMEM; BioWhittaker, Verviers, Belgium) supplemented with 10% fetal bovine serum (FCS; Gibco, Grand Island, NY, USA), 200 mM L-glutamine, penicillin (100 mg/ml) and streptomycin (100 mg/ml) and maintained at 37°C in 5% CO2. Cells were transfected using Lipofectamine 2000 (Invitrogen-Thermo Fisher Scientific) according to the manufacturer’s protocol. For immunofluorescence experiments, cells were washed in 1X PBS and then fixed for 20 min in 4% paraformaldehyde, 4% sucrose, dissolved in 60 mM PBS pH 7.4. After permeabilization in 0.25% Triton/PBS for 10 minutes, cells were incubated for 1h in blocking solution (1X PBS, 0.1% Tween 20, 3% BSA) and then incubated for 1h with the indicated primary antibodies in blocking solution in the humidified chamber at room temperature (RT). After three washes in PBS/0.1% Tween 20, coverslips were incubated with the secondary antibody for 1h at RT, washed three times in PBS/0.1% Tween and mounted with DAPI/VectaShield (Vector Laboratories) on microscope slides. For oxidative stress, cells were exposed from 0.5 to 1 mM sodium arsenide in complete medium for 30 min at 37°C. Cells were observed using a Zeiss Axioplan microscope equipped with a CCD camera (Photometrics, Tucson, AZ). Images were acquired using the ImageJ and processed with Adobe Photoshop.

### Larval brains immunofluorescence

Third-stage larval brains were dissected in physiological solution, fixed in PBS+3.7% formaldehyde for 20 minutes, briefly dipped for 30 seconds in 45% acetic acid (in ddH_2_O), and then transferred for 2 minutes onto a coverslip with a drop of 60% acetic acid (in ddH_2_O). Obtained preparations were squashed and flash-frozen in liquid nitrogen. The coverslip was removed, and the slides were immersed in ice cold absolute ethanol for 10 minutes and washed 2 times for 10 minutes in PBT (1X PBS with 0.1% Triton). Fixed preparations were incubated overnight with the indicated primary antibodies in PBS in a humid chamber at 4°C. The following day, the slides were washed twice in PBS for 5 minutes and then incubated for 1h at RT with secondary antibodies in PBS. After 2 additional washes in PBS for 5 minutes, the slides were mounted with DAPI/VectaShield and stored at –20°C. Cytological analysis was performed using a Zeiss Axioplan microscope equipped with a CCD camera (Photometrics, Tucson, AZ). Images were acquired using the ImageJ program and processed with Adobe Photoshop.

### Larval muscle immunofluorescence and confocal imaging

Larvae were dissected in ice-cold Ca^2+^-free HL3 saline and fixed in 4% formaldehyde for 10 min and washed in PBS containing 0.05% Triton X-100 (PBST) for 30 min. After washing, larval fillets were stained with phalloidin–TRITC (1:300 diluted in PBST, Sigma) for 40 min at RT and subsequently washed for 3x 20 min with 0,05% PBST. Larvae were mounted in DAPI/VectaShield. Confocal microscopy was performed with a Leica SP8 confocal microscope (Leica Microsystems, Germany). Confocal imaging of larval fillets was done using a z step of 0.5 μm. The following objective was used: 63× 1.4 NA oil immersion for confocal imaging. All confocal images were acquired using the LCS AF software (Leica, Germany). Images from fixed samples were taken from third instar larval fillets (segment A2, muscle 6/7).

### Larval fat bodies immunofluorescence

Larvae were placed in ice-cold PBS 1x, dissected in 4% formaldehyde, and fixed for 30 min. After fixation open larvae were washed in PBS containing 0.01% Triton X-100 (PBT) 2 x 10 min and blocked for 1h in Blocking Solution (BS) 3% BSA,1% NGS, PBT Then open larvae were incubated ON at 4° with anti-HA rabbit (1:250 in BS, Abcam) and anti-Fibrillarin rabbit (1:500 in BS, Gene Tex). The day after, open larvae where washed 2×10 min in PBS-Triton 0,01% and incubated 2h at RT with FITC-conjugated goat anti-rabbit (1:50 in PBT; Jackson Laboratories), AlexaFluor 555-conjugated donkey anti-mouse (1:50 in PBT; Invitrogen) and DAPI (1:300 in PBT). Open larvae were then rinsed in PBT, and the fat bodies isolated, placed on poly-L-lysine slide, and mounted in anti-fade mounting medium. Fat bodies preparations were analysed on a fluorescence microscope (Zeiss Apotome) and images acquisition performed using the Zen Pro software (Zeiss).

### DM2 derived cells immunofluorescence

Myoblast cells were seeded at a density of 1 × 10⁴ cells/cm² in CultureSlides chambers (Falcon, Cat# 354118, Corning Inc.). Cells were fixed with 4% paraformaldehyde and permeabilized using 0.5% Triton X-100. Following a 30-minute blocking step in 3% BSA, cells were incubated overnight at 4 °C with the primary antibody. The following day, cells were incubated with the appropriate fluorophore-conjugated secondary antibody and counterstained with Hoechst. Confocal images were acquired at 63× magnification using a LSM 980 confocal laser-scanning microscope equipped with Airyscan 2 (ZEISS, Germany).

### RNA extraction and quantitative PCR

Total mRNA was isolated from *Drosophila* larval fat bodies by using Trizol (15596026, Thermo Fisher Scientific) according to the manufacturer’s instructions. RNA was reverse-transcribed (1 mg each experimental point) by using SensiFAST cDNA Synthesis Kit (BIO-65053, Bioline) and qPCR was performed as described (18) using SensiFast Sybr Lo-Rox Mix (BIO-94020, Bioline). The run was performed by using the Applied Biosystems (Waltham, MA) Quant Studio 3 Real-Time PCR System 36 instrument (Marzullo et al., 2023b). Primer Sequences are reported in Resource Table.

### Image Analysis, Representation, and Statistical analyses

Pictures of fly eyes were taken with the stereomicroscope (ZEISS Semi 508, 50X) equipped with Axiocam camera 212 using ZEISS ZEN software (blue edition). Eye area was measured with the open-source software NIH ImageJ FIJI. Images of *Drosophila* larval fat bodies were analyzed using Fiji software. The number of cells with fragmented nucleoli was quantified and expressed as a proportion of the total number of cells per field. Fragmentation was defined based on nucleolar morphology and shape, as identified in the fibrillarin channel. Nucleolar size was measured by outlining individual nucleoli and calculating the corresponding area in Fiji.To quantify the number of puncta in each image, an arbitrary but fixed region of interest (ROI) of identical size was selected for all images. The analysis was performed three times per image, and the number and size of puncta within each ROI were quantified using the “Analyse Particles” function in Fiji (ImageJ). Co-localization analysis was performed by Fiji tolls using JACoP plug-in in order to obtain the Pearson coefficient from single ROIs of original images. The Pearson correlation coefficient measures the strength and direction of the linear relationship between two variables. Values between ±0.50 and ±1.00 indicate a strong correlation; values between ±0.30 and ±0.49 indicate a moderate correlation; and values below ±0.29 indicate a weak correlation. A value of zero implies no linear relationship. Statistical analysis was performed using Prism 10 software (MacKiev). The Shapiro-Wilk Test was used to assess the Normal distribution of every group of different genotypes. Statistical differences for multiple comparisons were analysed with the Kruskal-Wallis for non-parametric values or with one-way ANOVA for parametric values. The Dunn’s or the Tukey’s test was performed, respectively, as Post-Hoc Test to determine the significance between every single group. The Mann–Whitney *U* test or the t-test were used for two groups comparison of non-parametric or parametric values, respectively. Statistical significance was defined at P< 0.05 (∗), P< 0.01 (∗∗), P< 0.001 (∗∗∗), and P< 0.0001 (∗∗∗∗).

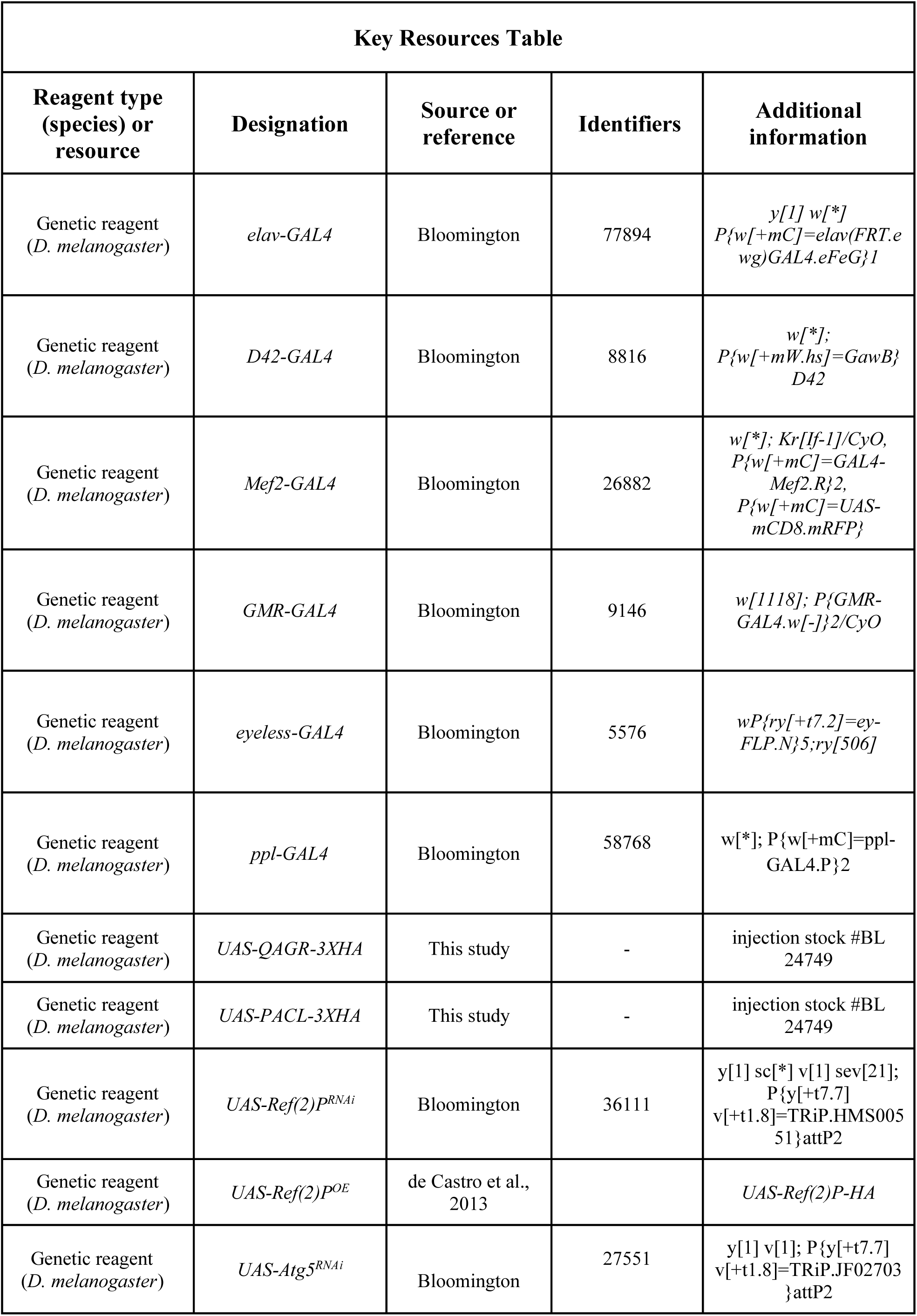
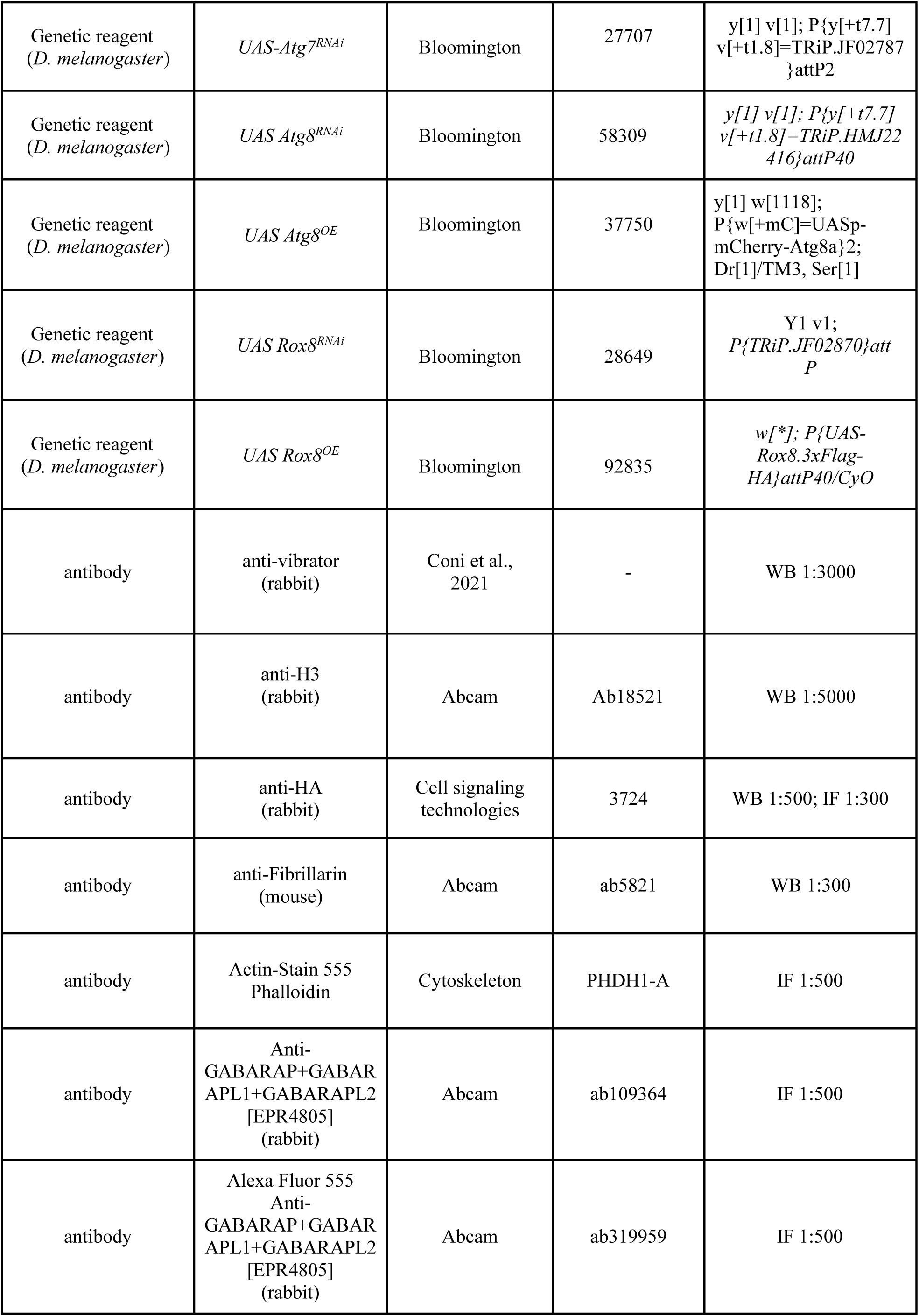
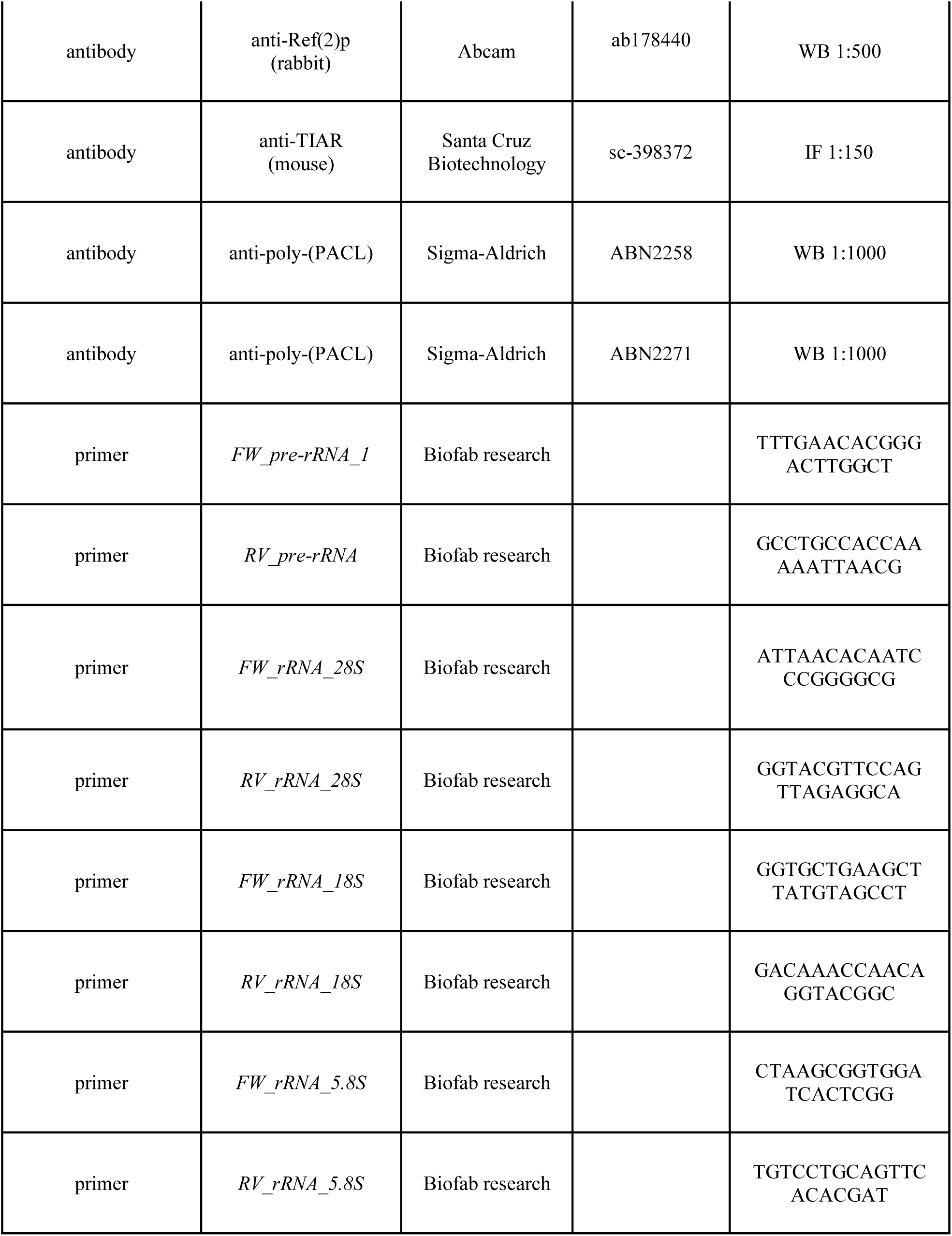
Key Resources Table.

## Acknowledgments

This work was supported by AFM-Telethon to MM (project number 28731), by the National Recovery and Resilience Plan (NRRP), Mission 4, Component 2, Investment 1.1, Call for tender No. 1409 published on 14.9.2022 by the Italian Ministry of University and Research (MUR), funded by the European Union – NextGenerationEU–CUP B53D23024800001-Grant Assignment Decree No. P2022S7H3H to LC and by Telethon to GC (project number 4398). We thank L.M. Martins for sharing the UAS-Ref(2)P-HA fly stock.

